# Active learning-guided mechanistic modeling of CXCL9 regulation in pancreatic cancer

**DOI:** 10.1101/2025.10.24.684332

**Authors:** B. Wang, V. S. M. K. Yelleswarapu, L. Descamps, F. Eduati

**Author notes:** corresponding author, Contact information, Lead contact: Eduati F.

## Abstract

Cold tumors like pancreatic cancer suffer from poor immune infiltration, limiting effective anti-tumor responses. The chemokine CXCL9 promotes immune cell recruitment, but the signaling mechanisms regulating its expression in tumor cells remain poorly understood and underexplored as targets for modulation. We present a framework that integrates active learning with mechanistic logic-ODE models to guide perturbation screenings and uncover regulators of CXCL9 in pancreatic cancer cells. Using perturbation-response data and curated prior knowledge, we trained interpretable models to identify signaling mechanisms that enhance CXCL9 expression and prioritize drug combinations. Active learning enabled data-efficient model refinement and guided informative experiments under resource constraints. Benchmarking on synthetic data and experimental validation confirmed the performance of different acquisition strategies and revealed cell line-specific regulatory differences. Our results provide insight into tumor cell-intrinsic control of CXCL9 and demonstrate how combining active learning with mechanistic modeling supports rational, targeted experimental design.

## Introduction

Pancreatic ductal adenocarcinoma (PDAC) is one of the most aggressive and lethal malignancies, characterized by late diagnosis, rapid progression, and limited treatment options ^1,2^. Despite the success of immune checkpoint blockade (ICB) therapies in several other cancer types, their efficacy in cold tumors like PDAC has been disappointingly low ^3^. This is largely attributed to PDAC’s profoundly immunosuppressive tumor microenvironment (TME), which is marked by low infiltration of cytotoxic T cells and a dense stroma that supports immune evasion^4^. Interestingly, rare PDAC cases exhibiting high CD8⁺ T-cell infiltration are associated with improved prognosis, highlighting the therapeutic potential of strategies that convert “cold” tumors into T-cell–inflamed phenotypes ^5,6^. In this context, the chemokine CXCL9 has drawn increasing attention as a key modulator of immune infiltration. CXCL9 recruits effector CD8⁺ T cells ^7^ and is associated with improved immunotherapy responses across multiple cancer types^8^. Deciphering how CXCL9 expression is regulated in pancreatic cancer cells could therefore uncover actionable mechanisms to modulate the TME and improve therapeutic outcomes.

To elucidate CXCL9 regulatory mechanisms in PDAC cells, mechanistic modeling offers a valuable framework. Computational models created by combining prior knowledge of intracellular signaling pathways with experimental perturbation data can reveal causal regulatory interactions and predict the effects of targeted interventions ^9^. This approach better captures mechanistic insights compared to purely data-driven models, which focus primarily on relationships between perturbed and observed variables ^10^. Among available formalisms, logic-based ordinary differential equations (logic-ODEs) strike a balance between biological interpretability and computational tractability ^11^. Logic-ODEs extend qualitative logic rules into a continuous dynamic system, allowing for the simulation of graded responses, including feedback loops, while preserving the modularity and transparency of logic models ^10^. Crucially, they do not require detailed kinetic parameters, which are often unknown in cancer signaling networks, and can incorporate both literature-derived topology and context-specific experimental data ^12,13^. This makes them particularly suitable for modeling the regulation of specific outputs, such as CXCL9 expression, in response to diverse perturbations of signaling components.

Training mechanistic models such as logic-ODEs requires perturbation-response data that inform how the system reacts to targeted interventions. In practice, this involves exposing cells to various stimuli or inhibitors and measuring downstream effects, such as CXCL9 expression. However, as signaling networks grow in size and complexity, the number of possible single and combinatorial perturbations increases exponentially, making exhaustive experimental testing unfeasible. This challenge is particularly significant when sample material is limited or when readouts are costly. Active learning provides a principled framework to address this issue by iteratively selecting the most informative experiments. In each round, new perturbations are chosen to maximize expected model improvement, allowing more efficient exploration of the experimental design space and better model generalization ^14,15^. This hypothesis-driven, iterative approach has been used increasingly for biomedical applications such as compound or antibody screenings to streamline effective drug discovery ^14,16,17^. An important reason for this trend is that previous studies have demonstrated that combining active learning with machine learning models can improve performance in biological applications. In some cases, models trained with active learning outperform those trained on larger but randomly selected datasets ^15,18^. Despite this progress, active learning has not been applied in combination with mechanistic biological models. This is partly due to computational challenges and the lack of flexible software frameworks. Instead, recent efforts have focused on building surrogate models that approximate the behavior of mechanistic simulations ^19^. While surrogates enable active learning in principle, they disconnect the experimental design process from the original biological model and may introduce approximation errors. By contrast, directly coupling active learning with mechanistic models allows experimental design to remain grounded in prior biological knowledge, potentially leading to more interpretable and robust discoveries.

In this study, we present an integrated experimental and computational pipeline to investigate the regulation of CXCL9 expression in pancreatic cancer cells. Starting from a prior knowledge signaling network, we trained context-specific logic-ODE models using perturbation-response data collected from two PDAC cell lines. These models enabled us to analyze the signaling mechanisms underlying CXCL9 regulation through parameter sensitivity analysis, *in silico* knockdowns, and bootstrapping. To extend the predictive power of these models and guide new experimental investigations, we implemented an active learning framework that directly interacts with the mechanistic model to prioritize informative perturbations. We first benchmarked multiple sampling strategies *in silico* and then used the most promising approach to predict drug combinations expected to enhance CXCL9 expression. Selected drug combinations were subsequently validated through additional follow-up wet lab screenings, demonstrating the utility of the pipeline in identifying effective treatments that modulate immune-relevant pathways.

While our focus in this study was on CXCL9, the proposed pipeline can be broadly applied to different mechanistic models of intracellular signaling to support data-efficient model construction, analysis, and experimental design.

## Results

### Prior knowledge network curation and experimental design

We selected AsPC1 and BxPC3, two pancreatic cancer cell lines with distinct drug response profiles based on our reanalysis of the Genomics of Drug Sensitivity in Cancer (GDSC) dataset^20^. AsPC1 showed lower overall sensitivity and poor correlation with BxPC3 responses (median Z-scores: 2.53 vs. 0.2; Spearman r = 0.19, p = 0.0018), suggesting the need for cell line-specific models. (**Supplementary Fig. 1**).

To investigate upstream regulation of CXCL9, we reviewed literature studies and reanalyzed public databases to identify relevant proteins and signaling pathways. Two cytokines are well-established CXCL9 inducers: IFNγ and TNFα ^21^. These cytokines activate the signaling pathways JAK-STAT and NFKB, which are central to immune and inflammatory signaling (**Figure 1A**; ^21^). Both signaling pathways are also involved in complex crosstalk with other pathways such as PI3K-AKT, MAPK and p53. Therefore, in addition to IFNγ and TNFα stimulation, we included inhibitors targeting key components of these pathways: momelotinib (JAKi), BMS-345541 (IKKi), taselisib (PI3Ki), trametinib (MEKi), and salirasib (RASi). While JAKi and IKKi directly modulate the major CXCL9 regulators (JAK-STAT, NFKB), analysing published transcriptomic data we observed that PI3Ki and MEKi potentially affect CXCL9 expression (**Supplementary Fig. 1C-D**) ^22^. Additionally, RASi, though of clinical interest in pancreatic cancer, has an uncertain role in CXCL9 regulation ^23^. Altogether, pairwise combinations of the two cytokines and five inhibitors were used to generate perturbation-response data for model training (**Figure 1A**).

**Figure 1.**
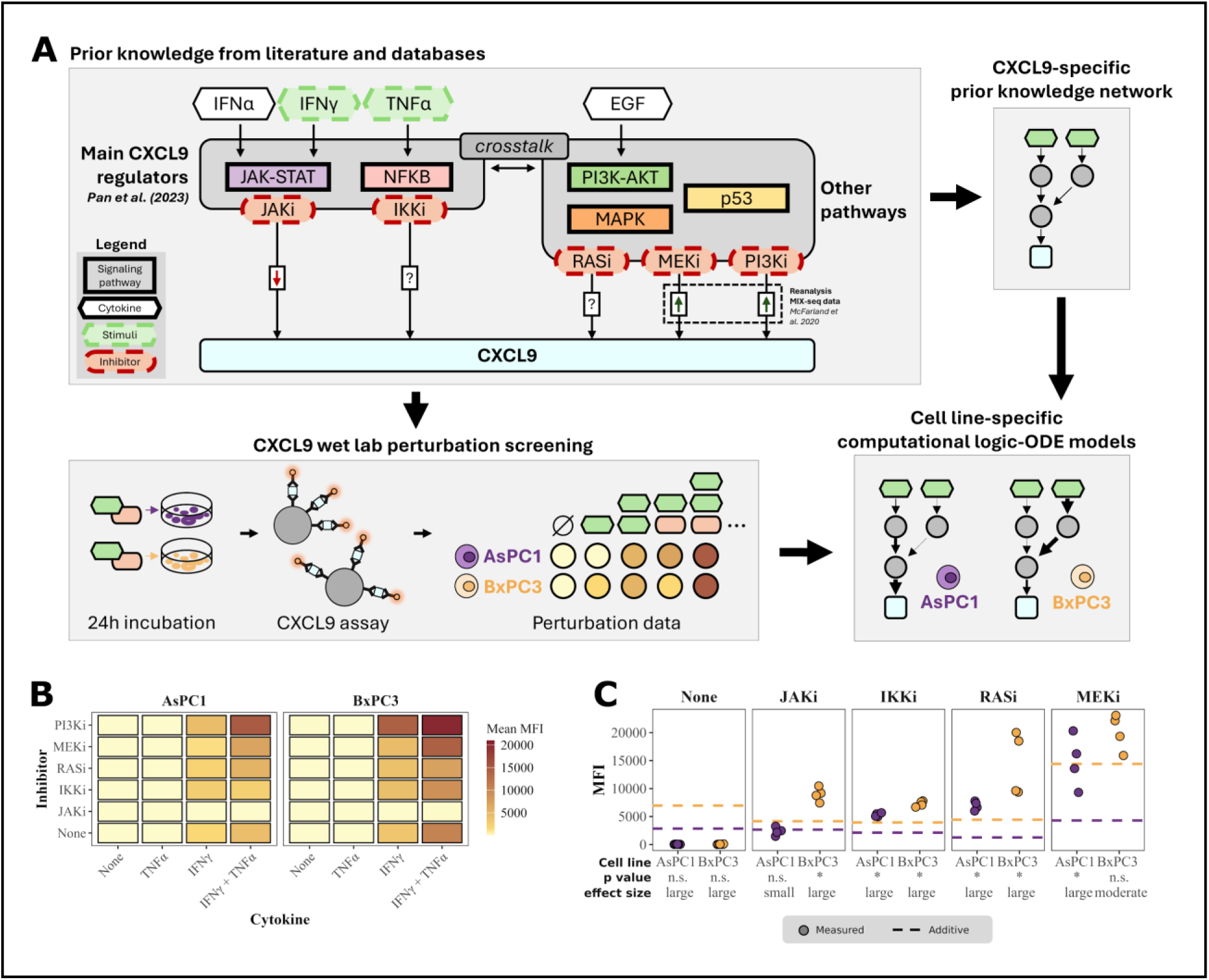
Prior knowledge network (PKN) and perturbation screenings. **(A)** Overview of input preparation for CXCL9 logic-ODE model. Prior knowledge was used to set up the perturbation screening and prior knowledge network, also giving an idea of what possible effects could be of inhibitors (red arrow = inhibitory, green arrow = stimulatory, question mark = unknown) **(B)** Measured CXCL9 levels from wet lab screenings, reported as Mean Fluorescent Intensity (MFI), averaged across two independent screenings with duplicates. **(C)** Statistical analysis of IFNγ and TNFα synergy. A Wilcoxon rank sum test compared measured values of the IFNγ + TNFα condition from the wet lab with additive values, calculated as the sum of CXCL9 responses from IFNγ and TNFα only conditions.

Moreover, to model CXCL9 regulation, we used the gathered knowledge on CXCL9 regulation to construct a prior knowledge network (PKN) (**Figure 1A**). We included relevant transcription factors for CXCL9 regulation identified using literature and DoRothEA ^24^. Transcription factors were linked to their upstream cytokine inducers (EGF, IFNα, IFNγ, TNFα) to create a CXCL9-specific PKN (**Supplementary Note**). In this network, nodes represent proteins and edges are directed interactions which can be either stimulatory or inhibitory. AND gates are a special type of node, which only affect nodes downstream if all upstream regulators are active.

### Shared and distinct CXCL9 responses in AsPC1 and BxPC3 cell lines

We quantified CXCL9 protein levels using a bead-based sandwich immunoassay with flow cytometry readout, across two independent wet lab screenings with duplicates per cell line (see **Methods**). AsPC1 and BxPC3 displayed similar patterns of CXCL9 regulation in response to perturbations, with some notable cell line–specific differences (**Figure 1B, Supplementary Fig. 2A**). BxPC3 consistently showed higher CXCL9 expression than AsPC1. As expected, in both cell lines IFNγ alone or combined with TNFα significantly upregulated CXCL9, while JAKi strongly suppressed CXCL9, confirming the key role of JAK-STAT signaling ^21,25,26^. Interestingly, PI3Ki and MEKi increased CXCL9 expression, especially when combined with the stimulation of both cytokines. In contrast, IKKi suppressed CXCL9 expression in AsPC1, but only under dual cytokine stimulation, and had minimal effect in BxPC3. RASi had negligible effects on CXCL9 in either cell lines, consistent with limited known involvement in CXCL9 regulation ^23^.

From the wet lab screenings, we observed minimal induction of CXCL9 in response to TNFα alone. However, when combined with IFNγ, the effect exceeded that of IFNγ alone, suggesting potential synergy between the two cytokines. To test this, we performed a Wilcoxon rank sum test to assess whether the combined cytokine effect significantly exceeded the sum of the individual effects ^27,28^ (**Figure 1D**). Synergy was confirmed for both cell lines in absence of inhibitors or with RASi or MEKi (p = 0.03, Cohen’s d = 0.82, for both AsPC1 and BxPC3).

Interestingly, PI3Ki disrupted synergy in BxPC3 (p = 0.47, d = 0.31) but not in AsPC1 (p = 0.03, d = 0.82), while IKKi disrupted synergy in AsPC1 (p = 0.89, d = 0.10) but not in BxPC3 (p = 0.03, d = 0.82). These findings support distinct cell line–specific responses, especially to IKKi, despite overall similarities in CXCL9 regulation.

### *In silico* knockouts and sensitivity analysis identify key regulators and cell line–specific control of CXCL9

Next, using CellNOptR (Terfve et al. 2012) we converted our PKN with into a set of logic-based ordinary differential equations (logic-ODEs) and optimized cell line–specific models for AsPC1 and BxPC3 by minimising the mean squared difference between the model prediction and the measured CXCL9 responses from the wet lab screenings. For each cell line, we trained an ensemble of ten models, which showed strong agreement with the experimental data (Pearson correlation, R² = 0.998 for AsPC1, 0.995 for BxPC3, p < 0.01; **Supplementary** Fig. 2B), confirming the models’ ability to capture the observed perturbation responses.

Importantly, model parameters reflect the strength of regulatory interactions, making them biologically interpretable. To investigate the regulatory mechanisms underlying CXCL9 expression, we performed *in silico* single-edge knockouts using our trained logic-ODE models. Most knockouts had limited effect, but several interactions within the NFKB and JAK-STAT pathways had notable impact in one or both cell lines (Figure 2A-B). Specifically, ERK-AR knockout reduced CXCL9 expression in BxPC3 (p = 0.004), while JAK-STAT1 and STAT1-CXCL9 knockouts affected only AsPC1 (p = 0.016 and p = 0.017, respectively). We also tested how individual knockouts influenced the synergy between IFNγ and TNFα. Interactions in the NFKB pathway were the most disruptive in both cell lines, while JAK-STAT interactions— specifically IFNGR-JAK, JAK-STAT3, and STAT3-AR—disrupted synergy only in BxPC3 (Figure 2C), suggesting a more prominent role of JAK-STAT signaling in mediating synergy in this cell line.

**Figure 2.**
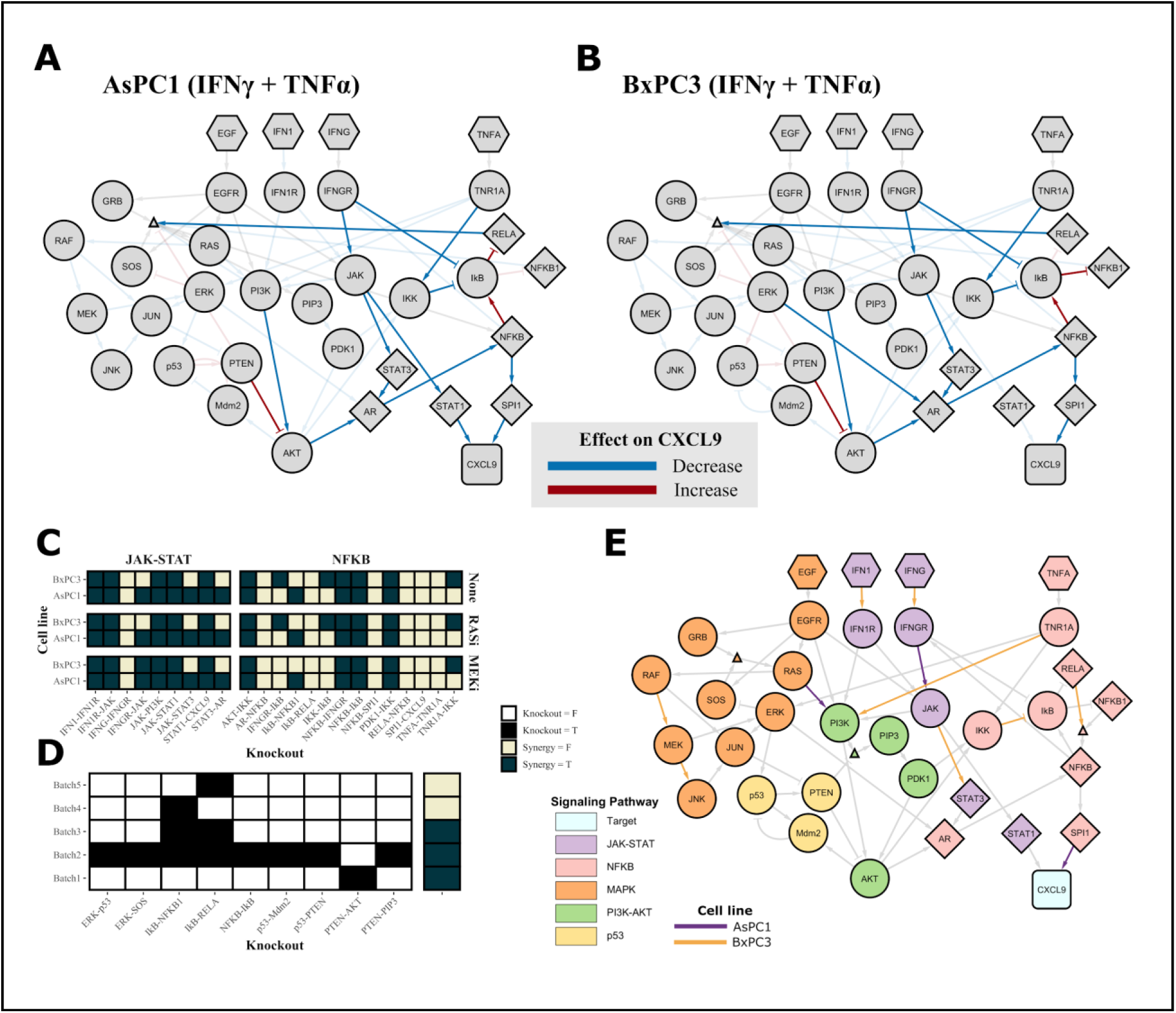
Parameter analysis of AsPC1- and BxPC3-specific models. **(A-B)** Effects of *in silico* single-edge knockouts on CXCL9 responses under IFNγ + TNFα stimulation. Significant effects are highlighted separately per cell line, indicating whether CXCL9 increased or decreased. **(C)** Effect of *in silico* single-edge knockouts on IFNγ and TNFα synergy. Following each knockout, a significance test was performed to determine whether synergy was retained (T) or lost (F). In the original (non-knockout) models, synergy was present across all visualized conditions (no inhibitor, RASi, or MEKi). **(D)** Effect of *in silico* knockouts of different sets of edges (batches) on IFNγ and TNFα synergy. Synergy was assessed after removal of each batch. **(E)** Parameter comparison between AsPC1 and BxPC3 models using bootstrapped parameter values. Significant differences are highlighted in the network.

Building on these findings, we simulated combinatorial knockouts to identify potential interventions that could restore synergy when it was lost. In BxPC3, synergy was disrupted under PI3Ki), likely due to elevated CXCL9 levels in the IFNγ + PI3Ki condition. Multiple single-edge knockouts, particularly in the JAK-STAT and NFKB pathways, were sufficient to restore synergy by reducing CXCL9 levels (**Supplementary** Fig. 3). In contrast, upregulation of CXCL9 was more difficult to achieve. In AsPC1, synergy was lost upon IKKi, and only the PTEN-AKT knockout restored it at the single knockout level. We therefore tested combinations of edges that increased CXCL9 (excluding PTEN-AKT) and found that simultaneous knockouts of ERK-p53, ERK-SOS, IkB-NFKB1, IkB-RELA, NFKB-IkB, p53-Mdm2, p53-PTEN, and PTEN-PIP3 reestablished synergy (p = 0.014). Next, we decreased the number of knockouts and found that dual knockouts of IkB-NFKB1 and IkB-RELA restored synergy (p = 0.11), while individual knockouts had no significant effect (p = 0.97 and p = 0.47, respectively; Figure 2D). These results show how mechanistic models can be used to simulate regulatory interventions that rescue context-dependent loss of cytokine synergy.

To validate our knockout findings, we conducted a multi-parametric sensitivity analysis (MPSA) and bootstrap resampling across 100 model fit per cell line. Key parameters within the JAK-STAT and NFKB pathways were consistently identified as major regulators of CXCL9 (**Supplementary** Fig. 4A). Most parameters were stable across bootstrap runs, except those in the NFKB pathway, which showed greater variability, likely due to its complex feedback loops involving IKK, IkB, and NFKB. Low correlations among NFKB-related parameters (Spearman r between −0.25 and 0.25; **Supplementary** Fig. 4B-C) suggest potential identifiability issues in this pathway. Finally, comparison of parameter distributions between cell lines showed general agreement, but also quantitative differences in pathway strength. Eight parameters (IFN1-IFN1R, RELA-NFKB, IKK-IkB, RAF-MEK, MEK-JNK, TNR1A-PI3K, IFNG-IFNGR, and JAK-STAT3) were significantly higher in BxPC3 (p < 0.05), while two (IFNGR-JAK, SPI1-CXCL9) were significantly higher in AsPC1 (p < 1.83E-4; Figure 2E). These results support both the interpretability and robustness of the models and provide mechanistic insights into cell line– specific regulation of CXCL9.

### An active learning pipeline to enable iterative model refinement and efficient experimental design

Next, we implemented an active learning pipeline directly coupled with our logic-ODE models, to efficiently explore the large space of possible perturbations and prioritize experimental testing of perturbations that influence CXCL9 expression and enable model refinement. The pipeline proceeds in iterative rounds: models trained on an initial set of perturbation-response data predict CXCL9 levels for all remaining candidate conditions. Based on these predictions, a subset of new conditions is selected using an acquisition function, experimentally (or *in silico*) tested, and added to the training set for model re-optimization. This cycle is repeated until no candidates remain or performance stabilizes (Figure 3, see **Methods** for more detailed description).

**Figure 3.**
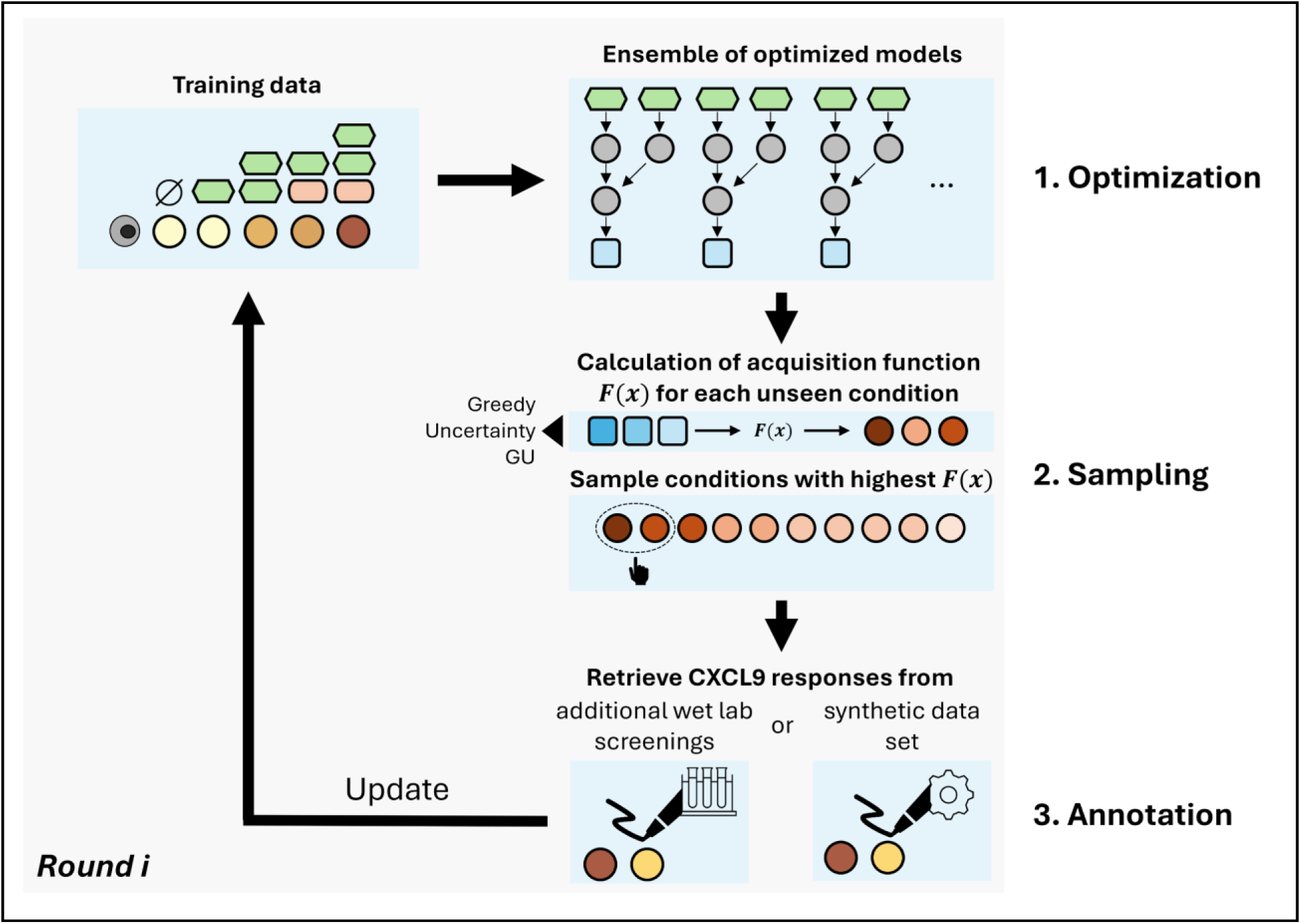
Active learning workflow. A schematic of the combination of active learning and logic-ODE models, including integration with data annotation from e.g. wet lab screenings

To evaluate the impact of key design choices in this workflow, we used a synthetic dataset generated from the trained AsPC1 model, with added Gaussian noise to mimic experimental uncertainty. We systematically tested the effect of different acquisition strategies, including greedy (which favors high predicted CXCL9), uncertainty (which prioritizes conditions with high prediction variance), greedy + uncertainty (GU) (which balances both criteria), and random sampling (see **Methods** for formal definition). We also evaluated different numbers of conditions added per round, ranging from 2 to 10 added conditions per round, and compared several initial training sets, including manual, random, and full selections. Model performance was assessed based on hit discovery and predictive power across unseen conditions. This benchmarking step enabled us to identify acquisition strategies and configurations with favorable trade-offs for downstream application to real perturbation data. The subsequent sections first present the *in silico* benchmarking results and then describe the application of the pipeline to *in vitro* perturbation data with experimental validation.

### Greedy sampling increases hit discovery, while uncertainty improves generalization

Using the synthetic dataset, we ran five rounds of active learning with ensembles of ten models to benchmark acquisition strategies. Greedy and GU acquisition functions identified more CXCL9-inducing conditions (“hits”) than uncertainty or random sampling (Figure 4A). For example, after five rounds, greedy and GU achieved 1.4 to 1.9 times more hits than random sampling, while uncertainty showed a more modest and less consistent advantage (1.0 to 1.6 more hits). However, greedy and GU also generated more false positives, especially at higher number of conditions added per round (N=6–10), as indicated by declining proportions of correctly predicted hits. In contrast, uncertainty maintained a more stable precision across different numbers of conditions added per round. These results suggest that adding a smaller number of conditions (N=2 or 4) is preferable for greedy and GU to reduce false positives, while uncertainty yields more reliable predictions even at higher number of conditions added per round.

**Figure 4.**
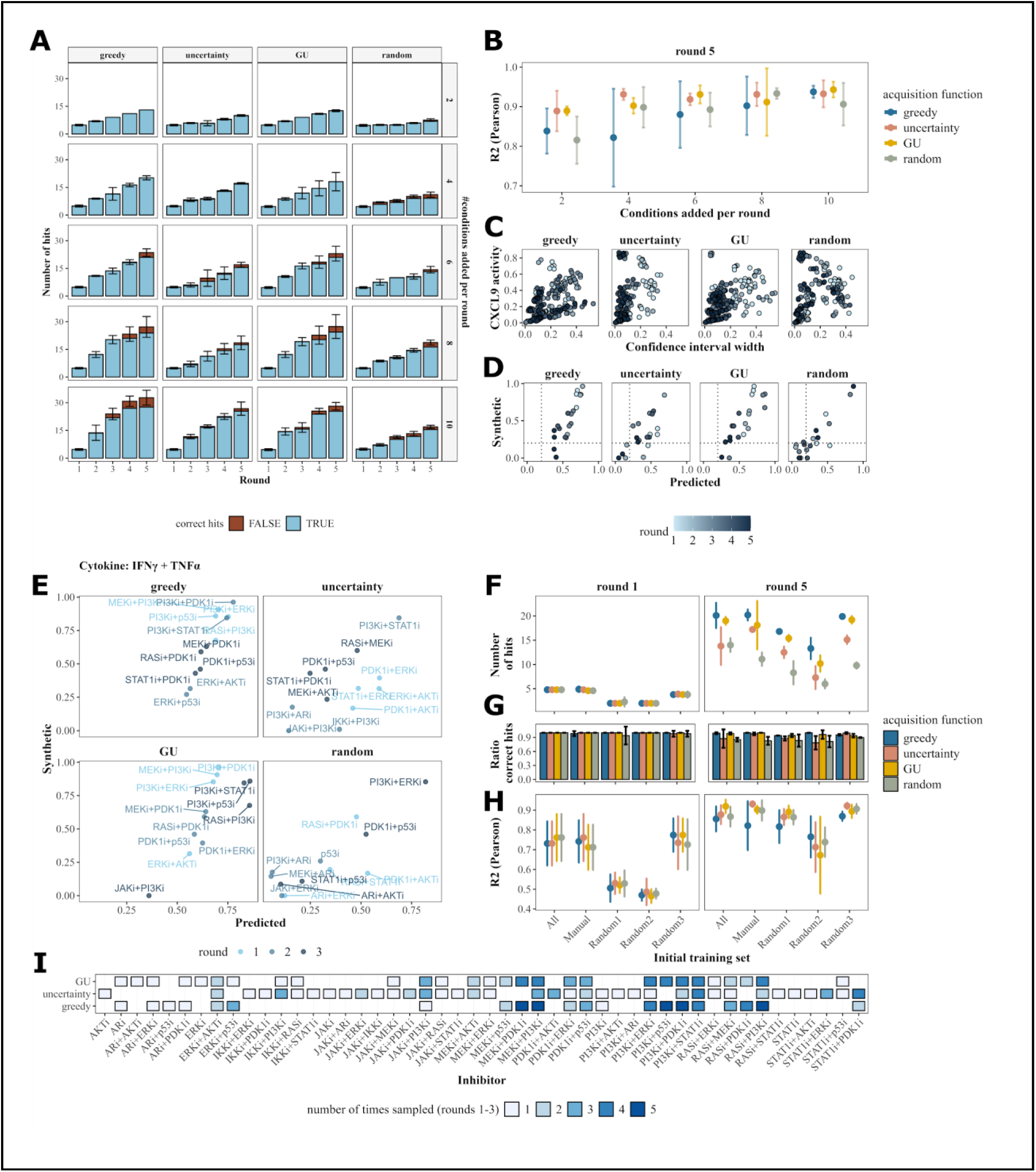
Active learning workflow with synthetic data. **(A-B)** Effect of the number of conditions added per round using different acquisition functions. Results are averaged across ten models per round; error bars represent standard deviation. **(A)** Hit discovery per round. From the total number of hits, colors indicate how many were correct versus incorrect. **(B)** Model generalization, represented by the Pearson correlation coefficient R^2^ after the final round (round 5). **(C-E)** In-depth characterization of models trained with four conditions added per round. **(C)** Comparison of mean CXCL9 activity and confidence interval width across acquisition functions. Each data point represents a condition from the selection pool in the corresponding round. **(D)** Comparison of model simulations with true CXCL9 values from the synthetic dataset. Each data point represents a condition added to the training set in that round. Dashed lines indicate hit boundaries, dividing the space into regions for true/false hits and non-hits. **(E)** Inhibitor combinations of added conditions during the first three rounds. **(F– H)** Effect of the initial training set on **(F)** hit discovery, **(G)** correct hit discovery, and **(H)** model generalization. Round 1 is compared with round 5 to illustrate how the starting set influences outcomes. **(I)** Frequency of selection for each condition characterized by one or two inhibitors across five different initial training sets (Manual, All, Random1, Random2, Random3)

Model generalization, measured as Pearson correlation (R^2^) between predicted and true CXCL9 levels from the synthetic data set across all conditions, including unseen ones, was consistently highest for uncertainty-based sampling. When adding four new conditions per round, uncertainty achieved the best overall R² (mean = 0.93), significantly outperforming all other acquisition strategies (p < 0.026, Figure 4B showing results for round 5). On the contrary, greedy showed lower generalization performances (mean = 0.88) and greater variability in R². Notably, R² values on the training set remained stable across all acquisition strategies and numbers of conditions added per round, indicating consistent fit to observed data (**Supplementary** Fig. 5A). These results emphasize that acquisition strategies strongly influence the performance of active learning with mechanistic models: greedy boosts hit discovery, uncertainty enhances generalization, and GU offers a valuable trade-off.

### Different acquisition strategies prioritize perturbations with distinct characteristics

To understand the mechanistic implications of acquisition function behavior, we examined which perturbation conditions were selected during active learning when four new conditions were added per round. We observed distinct sampling behaviors for the different acquisition functions. For instance, conditions that were selected only at later iteration rounds (Figure 4C, darker colour), had relatively low mean CXCL9 activities for greedy and low confidence interval width (CI) for uncertainty. GU maintained a balance, with values in later rounds tending to be low in both mean CXCL9 activity and CI. In contrast, random sampling did not show such trends, highlighting the advantage of acquisition functions in targeted sampling. Notably, uncertainty consistently yielded lower CIs than greedy and GU, reflecting its objective of reducing estimation variance. Additionally, comparing predicted vs. true CXCL9 responses highlighted a bias toward false positives over false negatives, as all acquisition functions, except for random sampling, selected two false positives and zero false negatives (Figure 4D).

Uncertainty showed great potential in correctly predicting also “non-hits” (i.e. conditions with low CXCL9 expression), while greedy and GU were biased toward high-response predictions. These results suggest that greedy is effective for prioritization of potential hits, but introduces estimation bias for true positive hits, whereas uncertainty offers a better balance between true positive and negative hits.

Next, we examined the selected drug combinations (Figure 4E). As expected, acquisition functions prioritize different sets of conditions. Greedy favored combinations containing PI3Ki— which we previously identified as strong CXCL9 inducers (Figure 1A-B)—while uncertainty focused on ERKi and AKTi combinations whose effects on CXCL9 remain uncharacterized. GU blended both approaches. The distinct sampling patterns help discover valuable candidates for further wet lab validation.

### Initial training set composition influences active learning outcome

In the preceding sections, we used an initial training set consisting of ten conditions manually selected to cover a broad range of perturbations affecting CXCL9 responses based on our experimental data (referred to as ‘Manual’, see **Methods**). To assess how the choice of initial training set influences hit discovery and model performance, we created four additional initial training sets (**Supplementary** Fig. 5B): one including all 24 conditions from wet lab screenings (‘All’), and three randomly selected sets of ten conditions (‘Random1’, ‘Random2’, ‘Random3’).

In round 1, Random1 and Random2 had fewer predicted hits, while Manual and All and Random3 performed similarly (Figure 4F). Across all rounds, the greedy approach consistently identified the most hits, followed by GU, uncertainty, and random sampling. Final average hit counts were highest for Manual (17) and lowest for Random2 (9), indicating that the quality of initial sampling impacts final outcomes. Despite differences in hit counts, the ratio of correct hits remained above 0.78 for all initial training sets, with random sampling yielding the most false positives (Figure 4G). Uncertainty was the most sensitive to the initial training set, with its correct hits ratio dropping below 0.88 (Random1, Random2, All), while remaining ratios were significantly higher (>0.98) for the remaining initial training sets (Random3, Manual). Especially the poorer performance with the initial training set All is interesting, although this could be due to the complete experimental data set being an imbalanced data set with more non-hits than hits. These results for Uncertainty contrast with the acquisition function’a stability across varying numbers of added conditions per round (Figure 4A), suggesting greater dependence on initial data.

Model generalizability (R²) also varied depending on the initial training set. In round 1, Random1 and Random2 showed poor model performance (R² < 0.53; Figure 4H), while Random3 (R² = 0.75) showed performance comparable with All (R² = 0.75) and Manual (R² = 0.73). After five rounds, Random1 recovered to match other sets (R² = 0.86), but Random2 remained the weakest (R² = 0.72), likely due to lacking key cytokine-containing conditions. GU and uncertainty showed the least variation across initial training sets (except in Random2), with either one being the best performing model depending on the initial set, reflecting their respective strengths in generalization. We also examined the drug combinations selected by different initial sets. Greedy and GU showed consistent sampling patterns, always selecting several key PI3Ki- and MEKi-containing combinations across all setups (MEKi with PDK1i/PI3Ki and PI3Ki with p53i/PDK1i/RASi; Figure 4I). In contrast, the uncertainty approach led to more variable selections, focusing on different areas of the search space.

Overall, while the choice of initial training sets do influence early model performance, particularly for uncertainty, the active learning pipeline can recover from suboptimal starting conditions. The Manual and All sets yielded comparable final performance, indicating that well-chosen small sets can be as effective as larger ones. Moreover, greedy and GU were more sensitive to the number of added conditions, but more stable across different initial training sets. These findings highlight how acquisition strategy and initial sampling interact, offering guidance for future experimental designs.

### Wet lab experiments validate differences between acquisition strategies

After benchmarking our active learning pipeline *in silico*, we next applied the workflow to real experimental data by returning to the lab to measure CXCL9 responses for conditions selected through active learning (Figure 5). Starting from the Manual initial training set, we performed two rounds of active learning for both AsPC1 and BxPC3 cell lines. In each round, ensembles of ten models per acquisition strategy (greedy, uncertainty, GU, and random) were trained and used to select four new conditions, which were experimentally tested for CXCL9 expression.

**Figure 5.**
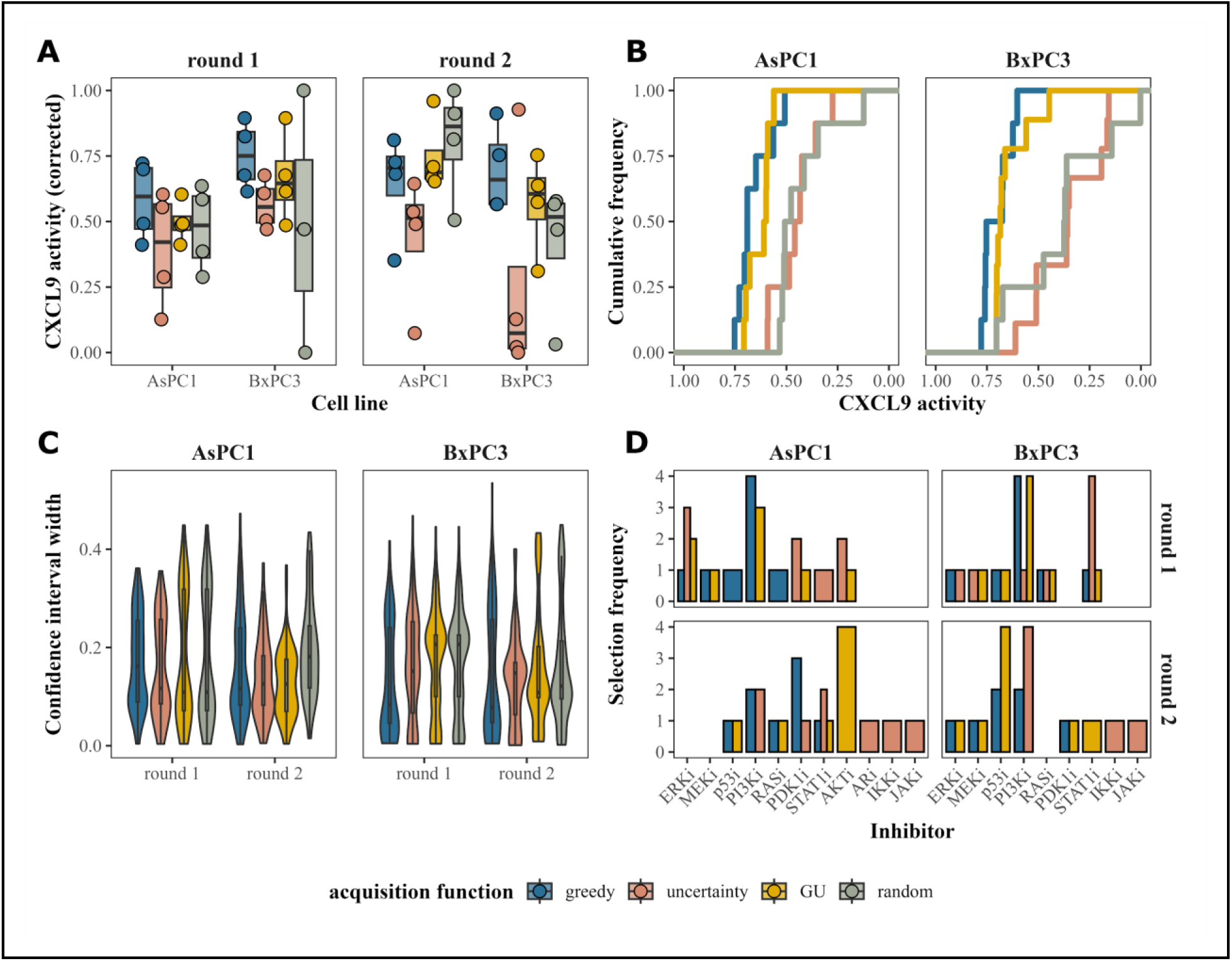
Integration of wet lab screenings in active learning workflow. **(A)** CXCL9 activities measured in AsPC1 and BxPC3 wet lab screenings for conditions selected by different acquisition functions. Averages were calculated from duplicate measurements. **(B)** Cumulative distribution functions of CXCL9 activity for each cell line and acquisition function. An early rise indicates that selected conditions yielded higher CXCL9 activity values. Greater distances between curves reflect larger differences in measured activity. **(C)** Confidence interval widths from models trained on conditions sampled in round 1 and 2. Violin plots show the distribution of widths from model simulations on all conditions in the selection pool. **(D)** Frequency of inhibitor selection, either as single agents or in combinations, across rounds and acquisition functions. A value of four indicates that the inhibitor was chosen by all acquisition strategies.

The selected perturbations primarily involved IFNγ and TNFα stimulation combined with inhibitors targeting ERK, AKT, PDK1, STAT1, AR, and p53. CXCL9 levels were quantified via immunoassays and reported as normalized mean fluorescent intensity (MFI), consistent with previous experiments (Figure 5A, see **Supplementary** Fig. 6 for perturbations per condition).

Surprisingly, several combinations, particularly those selected by greedy, resulted in lower-than-expected CXCL9 activity. For example, PI3Ki + ERKi, showed this effect in both cell lines. We hypothesized that higher cumulative drug concentrations in combination treatments (5 μM per drug) increased cell death, thereby reducing the number of viable cells capable of secreting CXCL9. In this scenario, individual surviving cells may still express high levels of CXCL9, but the overall secretion measured in the assay is lower due to reduced cell viability.. To test this, we performed a Caspase-3 (Cas3) assay to assess apoptosis levels. Strong negative Spearman correlations between Cas3 and CXCL9 were observed for conditions selected using the greedy approach (R = –1 and –0.8 in AsPC1 and BxPC3 respectively; **Supplementary** Fig. 6), while conditions selected using uncertainty had lower Cas3 values, suggesting less apoptosis.

As our current models do not account for apoptosis effects, we integrated confounding effects of apoptosis through combining normalized CXCL9 and Cas3 values, rescaling them to reflect CXCL9 secretion adjusted for apoptosis (**Methods**). Using the rescaled CXCL9 values, greedy-selected combinations showed the highest corrected CXCL9 levels, while uncertainty-selected combinations had the lowest. GU-selected conditions were intermediate, and random sampling showed more variability across conditions (Figure 5A). This ordering aligns with the *in silico* benchmarking, where greedy and GU prioritized higher-activity conditions, while uncertainty sampling favored conditions with greater model uncertainty, often resulting in more moderate measured responses. These findings underscore the reliability of the active learning framework and its potential to guide targeted experimentation.

In round 2, rankings shifted slightly, most notably with increased hits by random sampling in AsPC1 (Figure 5A). Still, greedy and GU consistently selected higher-activity combinations. Cumulative frequency plots confirmed this trend, greedy and GU sampled a greater number of conditions with high CXCL9 expression (defined as normalized CXCL9 > 0.6), especially in BxPC3 (Figure 5B). In contrast, uncertainty and random sampling explored a broader range of responses, including lower-activity conditions. The larger difference in CXCL9 activity between acquisition functions in BxPC3 compared to AsPC1 could be due to the cell line being more sensitive to perturbations resulting in a larger selection pool of possible hits.

While uncertainty is not expected to perform as well as greedy or GU in identifying strong CXCL9 inducers, as shown *in silico* and validated experimentally, it may offer advantages in improving model refinement. To evaluate this, we examined prediction confidence across all conditions over the active learning rounds (Figure 5C). Indeed, uncertainty sampling showed the most consistent reduction in prediction uncertainty, reflected in narrower and more stable confidence interval width across conditions. Although greedy had on average lower confidence intervals, it also showed greater spread, with several conditions having much higher uncertainty, likely due to its focus on a narrower region of the perturbation space. GU again showed intermediate behavior, balancing exploration and exploitation, and further supporting its value as a trade-off between hit discovery and model refinement. Nonetheless, we note that these trends are less pronounced than the differences observed in hit discovery and may require additional rounds to become more conclusive.

### Distinct selected targets reflect sampling strategy behavior

Beyond validating acquisition functions with experimental data, we examined which inhibitors were most frequently selected by each sampling strategy. Frequent selection, especially by greedy or GU, may indicate biological relevance. We tallied inhibitor appearances across rounds and cell lines and observed distinct patterns (Figure 5D). In round 1, PI3Ki was prominently selected by greedy and GU in both AsPC1 and BxPC3, appearing in every combination selected by greedy. Uncertainty showed no strong preference in AsPC1, but consistently selected STAT1i in BxPC3. Although STAT1i was expected to suppress CXCL9, screenings showed limited downregulation, possibly explaining why uncertainty prioritized it for further exploration.

By round 2, selections shifted and distinct patterns emerged between AsPC1 and BxPC3 (Figure 5D). PI3Ki remained prominent in BxPC3, but was favored by the uncertainty acquisition function. This result might reflect how PI3Ki’s induction of CXCL9 could, while not part of the main regulatory pathways, point to interesting targets within a less-characterized search space. GU frequently selected AKTi in AsPC1 and p53i in BxPC3, both of which were validated in the wet lab screenings as CXCL9 inducers (**Supplementary** Fig. 6). These selections may reflect either high uncertainty or expected CXCL9 activity, aligning with GU’s dual objective. Biologically, the emphasis on AKTi and p53i, both outside the canonical CXCL9 regulatory pathways, such as JAK-STAT or NFKB, suggests the involvement of less characterized signaling mechanisms. While p53 is a tumor suppressor and not an ideal inhibition target, its apparent role in CXCL9 induction in our dataset may point to indirect regulatory effects or interactions with other pathways. These observations could motivate further studies to identify alternative compounds that mimic this effect without compromising p53’s tumor-suppressive function.

## Discussion

Modulating the immunosuppressive tumor microenvironment (TME) remains a major challenge for improving the efficacy of immunotherapy in pancreatic cancer ^29,30^. To address this, we developed and validated a hybrid experimental–computational framework to systematically investigate how pancreatic tumor cells regulate secretion of chemokine CXCL9, an important driver of immune cell infiltration ^31^. By integrating data-driven perturbation screening with logic-ODE–based modeling and active learning, we prioritized combinations of signaling inhibitors that promote CXCL9 induction and provide mechanistic insights into context-specific regulatory interactions. This integrative approach provides a rational, systematic and interpretable way to guide perturbation experiments toward desired tumor cell-mediated immunomodulatory outcomes.

We started by curating a CXCL9-specific prior knowledge network (PKN) and performing cytokine and inhibitor screenings, leading to cell line-specific models. These models successfully captured experimental responses, such as CXCL9 upregulation by the PI3K inhibitor taselisib, and emphasizing the NFKB pathway’s central role in CXCL9 regulation. *In silico* knockouts revealed NFKB pathway’s involvement in cytokine synergy, particularly in AsPC1, suggesting that restoring NFKB activity may help recover synergistic cytokine response. This finding may explain the higher sensitivity of AsPC1 to the IKK inhibitor BMS-345541, as the IKKi is more effective in cells where NFKB signaling is pivotal ^32^. In addition, differences between AsPC1 and BxPC3 reflect previous findings linking aberrant NFKB signaling pathway behavior to drug resistance ^33,34^. These findings show how interpretable mechanistic models trained on targeted perturbation data can uncover cell-specific regulatory mechanisms that inform rational therapy design.

By coupling active learning with mechanistic modeling, we established a data-efficient strategy to identify informative perturbations under experimental constraints. Although this integration has rarely been explored due to the limited and context-specific nature of mechanistic datasets, our study shows that it is both feasible and beneficial. The curated prior knowledge network helped compensate for data scarcity by constraining the model hypothesis space, while the interpretable logic-ODE structure ensured that acquisition strategies remained mechanistically meaningful—greedy and GU focusing on CXCL9 induction, and uncertainty emphasizing model refinement. Together, these features enable interpretable, biologically grounded experimental design through active learning.

Our benchmarking experiments confirmed that active learning can be effectively applied to logic-ODE models. Simulations on synthetic data highlighted the importance of acquisition function choice, initial training set composition, and number of conditions added per round. Even when trained on sparse data, the models were able to capture combined effects of screened inhibitors (e.g., JAKi and PI3Ki) and generalize to unseen perturbations (e.g., ERK and p53 inhibitors), underscoring the ability of the framework to extrapolate from limited experimental conditions. Greedy and GU acquisition functions performed best for identifying CXCL9-inducing perturbations, while uncertainty-based sampling led to better generalization and model stability.

These findings were supported by *in vitro* validation experiments, which revealed biologically different responses to perturbations selected by each acquisition function. Conditions prioritized by greedy and GU led to stronger CXCL9 induction, while selected by uncertainty often targeted regulators with ambiguous or poorly constrained effects, offering insights into less-characterized parts of the network and helping to refine specific regulatory interactions.

Importantly, our framework supports iterative experimental design: model predictions can be continuously refined as new data become available, enabling adaptive experimentation. We observed that even when starting from small, manually selected training sets, the models rapidly converged toward high-performing perturbation conditions over successive rounds of active learning. This positions our approach as a scalable and data-efficient strategy for systematically exploring combinatorial perturbations in signaling networks.

This promising approach also highlights several opportunities for further development and improvement of the pipeline. First, model uncertainty is currently estimated from multiple model optimizations, but more principled approaches, for example, via Bayesian parameter inference such as BayesFit ^35^, could offer calibrated uncertainty estimates and support more nuanced acquisition decisions. Second, while our approach enables efficient exploration of combinatorial treatments, predictions are constrained by the structure of the prior knowledge network. The model can only simulate perturbations acting within the defined signaling axes. Expanding the PKN to include additional signaling axes, for instance, by incorporating insights from perturbation transcriptomics datasets such as MIX-Seq ^22^, could broaden the hypothesis space and allow active learning to operate over a more comprehensive set of potential regulatory mechanisms. Finally, while this study focused on a single immunomodulatory readout (CXCL9), future applications could benefit from multiplexed measurements — such as cell viability, apoptosis, or immunoinhibitory markers like PD-L1. Integrating such outputs into the active learning loop poses new challenges, especially when optimization goals differ across outputs (e.g., maximizing apoptosis while minimizing immunosuppressive signals), but offers a path toward richer and more holistic modeling of tumor–immune interactions.

Our data revealed a negative correlation between CXCL9 secretion and apoptosis, particularly in high-dose treatments, suggesting that cytotoxicity may reduce chemokine production by depleting viable, CXCL9-producing cells. This could help explain conflicting reports linking CXCL9 levels to both favorable and unfavorable prognosis in pancreatic cancer ^21,31,36^.

Incorporating cell viability markers into CXCL9 modeling could help disentangle these effects and better distinguish immunostimulatory from purely cytotoxic responses. To support this interpretation, we compared our results to drug sensitivity classifications from the GDSC database ^20^, which assesses compounds based on their ability to reduce cell viability.

Combining this with CXCL9 data enables identification of treatments with both cytotoxic and immunostimulatory effects. For example, AsPC1 cells were classified as insensitive to IKK inhibitors and showed reduced CXCL9 induction upon IKKi treatment, indicating limited benefit. In contrast, PI3K inhibition in BxPC3 moderately decreased viability while enhancing CXCL9 secretion, a more favorable dual effect. These examples highlight how integrating viability and immunomodulatory data can support more informed selection of perturbation strategies.

In summary, our framework combines mechanistic modeling with active learning to efficiently identify perturbation strategies that promote immunomodulatory responses in cancer cells. By integrating prior knowledge with experimental data, we demonstrate that interpretable, data-efficient modeling of chemokine regulation is feasible even in complex signaling contexts. The ability to iteratively refine model predictions based on targeted measurements opens new avenues for systematic exploration of treatment combinations. Beyond pancreatic cancer and CXCL9, the approach can be readily extended to other signaling outputs and cellular contexts, supporting rational, mechanism-driven design of combination therapies.

## Materials and methods

### Perturbation data

#### In-vitro CXCL9 screening

Two pancreatic cancer cell lines were used for the perturbation screenings: AsPC1 and BxPC3. These were cultured using RPMI-1640 medium supplemented with 10% FBS, 1% penicillin/streptomycin, L-glutamine and phenol red. We prepared 12-well plates for the CXCL9 assays containing RPMI-1640 medium with 1E6 cells seeded per microwell. The cytokines IFNγ and TNFα were used at 10 and 1 ng/mL for AsPC1 and BxPC3, respectively. Different cytokine concentrations were used due to the higher sensitivity of BxPC3 to CXCL9 inducement.

Inhibitors were selected based on two criteria: (1) inhibitor targets a protein that could be of interest potentially for CXCL9 regulation and (2) prior studies have demonstrated clinical relevance of the inhibitor for (pancreatic) cancer treatment, or showcased a response to the inhibitor. Alternatively, we also leveraged reanalysis of existing databases to identify possible inhibitors affecting CXCL9. Based on the first criteria, we selected five targets: JAK, IKK, RAS, MEK and PI3K. Next, we used the second criteria to select the following inhibitors: momelotinib (JAKi^37^), BMS-345541 (IKKi^32^), salirasib (RASi^23^), trametinib (MEKi^22^) and taselisib (PI3Ki^22^).

Inhibitors, diluted in DMSO, were added at a concentration of 5 μM. All samples in an inhibitor screening contained 1% of DMSO, which was not found to affect cell population numbers significantly. See **Table 1** for an overview of materials. A blank control is used in every screening and is defined as having no cytokines or inhibitors. Seeded cells were kept in an incubator (37 ⁰C, 5% CO₂) for 24h before retrieving the supernatant for the CXCL9 assay.

**Table 1:** Overview of materials and tools used.

| Reagents / tools | Reference / source | Identifier / catalog number |
| --- | --- | --- |
| Pancreatic cell lines |  |  |
| AsPC-1 | ATCC | CRL-1682 |
| BxPC-3 | ATCC | CRL-1687 |
| **Other reagents** |  |  |
| Recombinant Human CXCL9 (MIG) (carrier-free) | BioLegend | 578104 |
| LEGENDplex™ Human CXCL9 (MIG) Capture Bead B3, 13X | BioLegend | 740994 |
| LEGENDplex™ HU Proinflam. Chemokine Panel 1 Standard | BioLegend | 740987 |
| LEGENDplex™ HU Proinflam. Chemokine Panel 1 Detection Abs | BioLegend | 740986 |
| LEGENDplex™ Buffer Set A | BioLegend | 740368 |
| Invitrogen™ EnzChek™ Caspase-3 Activity Assay Kit | ThermoFisher Scientific | E13184 |
| Recombinant Human IFNγ (carrier-free) | BioLegend | 570204 |
| Recombinant Human TNFα (carrier-free) | BioLegend | 592702 |
| Recombinant Human IFNα2 (carrier-free) | BioLegend | 570102 |
| Momelotinib (JAKi) | MedChem Express | HY-10961 |
| BMS-345541 (IKKi) | MedChem Express | HY-10519 |
| Salirasib (RASi) | MedChem Express | HY-14754 |
| Trametinib (MEKi) | MedChem Express | HY-10999 |
| Taselisib (PI3Ki) | MedChem Express | HY-13898 |
| SCH772984 (ERKi) | Fisher Scientific | T6066 |
| Pifithrin (p53i) | Fisher Scientific | T2707 |
| MK-2206 (AKTi) | Fisher Scientific | T1952 |
| BX-795 (PDK1i) | Fisher Scientific | T1830 |
| Fludarabine (STAT1i) | Fisher Scientific | T1038 |
| Enzalutamide (ARi) | Fisher Scientific | T6002 |
| Dimethyl sulfoxide (DMSO) | Merck Life Science NV | 102952 |
| **Readout** |  |  |
| BD FACSSymphony A3 |  |  |
| BioTek Synergy H1<br>Microplate Reader |  |  |

We used the LEGENDplex™ HU Proinflam. Chemokine Panel 1 Standard Kit from BioLegend containing Human CXCL9 (MIG) Capture Beads B3, 13X (**Table 1**). This kit measures CXCL9 response through the formation of a sandwich assay complex consisting of the analyte, beads pre-coated with antibodies (diluted to 1X), biotinylated antibodies and streptavidin-phycoerythrin (SA-PE). We kept a 1:1 ratio between all assay components. The PE fluorescence was measured with the FACS SymphonyA3 to quantify CXCL9. The assay was performed on a 96-well plate according to the LEGENDplex™ protocol given for this kit.

#### CXCL9 data processing and analysis

We performed gating on the raw flow cytometry data to select the capture beads population, which were distinctive due to their intrinsic APC fluorescence. To remove outliers from this selected population, a lower limit was defined using Q1 - 1.5IQR and an upper limit using Q3 + 1.5IQR. Q1 and Q3 refer to the 25th and 75th quartiles, respectively. The IQR represents the interquartile range of the data. The Mean Fluorescent Intensity (MFI) was calculated as the arithmetic mean. We used normalization to scale the data to a range between 0 and 1, which is a requirement for the logic models. The condition with the lowest MFI was set as the minimum (0) and the condition with the highest MFI as the maximum (1). However, to avoid saturation, we capped the highest training value at 0.8 instead of 1. With such adjustment, the model can simulate higher CXCL9 levels than observed in the screenings. For statistical analysis, post-hoc pairwise comparisons with the Wilcoxon rank sum test were performed. We corrected for multiple hypothesis testing using the Benjamini-Hochberg Procedure. These non-parametric tests were chosen due to their high robustness against non-normality. Additionally, Cohen’s d effect size was calculated as the standardized mean difference between two groups. We selected a significance threshold of 0.05 (95% confidence interval).

#### Apoptosis assay

We measured the apoptosis marker Caspase-3 with the Invitrogen™ EnzChek™ Caspase-3 Activity Assay Kit from ThermoFisher, which includes cell lysis and quantification of Caspase-3 using Z-DEVD-R110 substrates. The fluorescent signals emitted by these substrates upon cleavage by Caspase-3 is measured with the BioTek Synergy H1 Microplate Reader set at measuring fluorescence at excitation/emission ∼496/520 nm. Preparation of the samples in a 96-well plate is performed according to the protocol supplied for this kit. We corrected for apoptosis by summing the normalized CXCL9 and Cas3 assay measurements (per assay), and rescaling these values to a range of 0-1.

### Logic-ODE model

#### Model structure

We used the logic-ODE formalism implemented in the CNORode add-on of CellNOptR ^10,11,13^. This approach requires two inputs: a scaffold model and experimental data. We curated a CXCL9-specific prior knowledge network (PKN) for the scaffold model. Nodes represent proteins and edges represent protein interactions. The network contains directional edges with a distinction between stimulatory and inhibitory protein interactions. The experimental data, also referred to as training data, were obtained through performing perturbation screenings measuring CXCL9 responses under different conditions. The MIDAS format, which is required for CellNOptR, encodes the experimental set-up using binary values ^38^. This format indicates presence (1) or absence (0) of stimuli (e.g. cytokines) and targeted drugs. These inputs force the activity of a node to 1 if it is a stimulus or to 0 if it is a targeted molecular inhibitor. Each condition has a unique combination of perturbations.

In the logic-ODE formalism, each node (*x_i_*) in the logic network is represented with an ODE (**Equation 1**) where *B_i_* is a continuous update function that indicates the regulation by *N_i_* upstream nodes. The sigmoidal transfer function *f*(*x_ij_*) (**Equation 2**) contains several tunable parameters: τ_i_ is the life-time of node *i, n_ij_* the Hill coefficient of the interaction between nodes *i* and *j*, and *k_ij_* is the strength of the regulation of node *j* on *i*. If, *j*=*k_ij_* = 0, no regulation is present and the higher the value, the stronger the regulation. For this study, *n* was fixed at 3, while τ and *k* were bounded between 0 and 1 ^11,13^. The initial activities of simulated proteins were set at 0.5 to allow for both up and down regulation, except for CXCL9 which was set to 0 as no downregulation was observed experimentally.

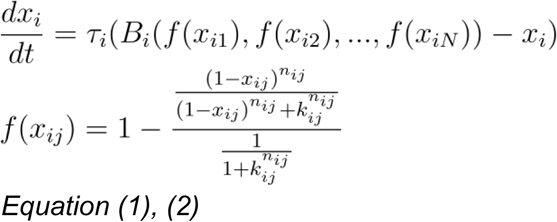

The CellNOptR workflow consists of the following steps: **(step 1)** import PKN and training data, **(step 2)** process network and **(step 3)** model training. In **(step 2)**, the complexity of the given logic model is reduced through compression while still retaining logical consistency ^39^. In **(step 3)**, model optimization is performed using the toolbox MEIGO with the metaheuristic enhanced Scatter Search. Advantages of this stochastic global optimization method include its robustness and ability to tackle complex optimization problems ^35^. The loss function for model optimization calculates the mean squared deviation between model prediction and training data and includes L1 regularization for the *k* parameter ^10,13^. The L1 regularization term has weight factor λ and sums the absolute values of every optimized *k* parameter. λ =1e-10 was used during optimization. Additionally, each model is required to have a best objective function value below 0.1 to ensure proper convergence.

#### Prior knowledge network

We curated a CXCL9-specific PKN to act as a scaffold network for CellNOptR. This PKN was curated via the following steps: **(1)** determining relevant transcription factors (TF) of CXCL9 that have a direct binding (i.e. TF binds to DNA site on gene) and functional binding (i.e. binding induces protein up-/downregulation) to the CXCL9 gene, **(2)** determining cytokines that affect CXCL9 levels, and **(3)** connecting TF and cytokines through established signaling pathways.

The resulting PKN was validated through a qualitative analysis, where we tested if simple effects from literature can be simulated correctly. See **Supplementary Note** for a more detailed description of PKN curation and validation. The final network consists of proteins from five major signaling pathways: JAK-STAT, NFKB, PI3K-AKT, MAPK and p53.

#### Parameter analysis

We compared parameter values between cell line-specific models using parameter distributions retrieved through bootstrapping, i.e. repeated optimization using sampled data with replacement^40^. We performed pairwise comparisons of the *k* parameter distributions from the two cell line-specific models. This was done with the Wilcoxon rank sum test and a significance threshold of 0.05. Additionally, we performed two types of parameter analyses: a global sensitivity analysis and *in silico* knockout experiments.

We implemented multi-parametric sensitivity analysis (MPSA) as described by Zi ^41^. Our goal was to retrieve high-level insights into sensitive model parameters for CXCL9. Latin hypercube sampling is used to generate different model parameter sets based on a parameter set from an optimized model. We did not fix any *k* parameter values and varied all of them for each parameter set. The resulting sum of squared differences (SSD) between CXCL9 model outputs and training data was compared with the SSD of the original parameter set. The differences were used to calculate the Kolmogorov-Smirnov (KS) statistic. The higher this value, which ranges between 0 and 1, the more impact the variation of a parameter has on the model output. The mean of dummy parameters is used to create a threshold between significant and non-significant KS statistics. Furthermore, we analyzed protein interactions and their relevance to CXCL9 responses with in-silico knockouts. Knockouts were mimicked in the model by setting the *k* parameter of an edge to 0. We compared simulations of the adjusted model with the original model. The bigger the difference, the bigger the impact of the knocked out interaction on the CXCL9 response.

### Active learning workflow

We developed an active learning pipeline to select the new conditions to test in the wet lab. Within the active learning loop, a set of trained models predict CXCL9 responses for unseen conditions in the candidate set. A subset of conditions is selected based on an acquisition function. After retrieving measurements from the wet lab, these conditions are added to the training set and removed from the candidate set. The previously trained models are further optimized on the updated training set. We define the moment in which the data sets change as the initialization of a new round within the active learning process. The loop continues with the next round until the candidate set is empty or if an early stopping criteria has been met.

#### Sampling

In each round of the active learning workflow, samples from the candidate set are selected by ranking conditions based on scores computed with an acquisition function. This function takes the predictions of an ensemble of models into account. Different types of acquisition functions are used, based on prior work by Vasanthakumari et al. (2024) who investigated active learning strategies for anti-cancer drug screenings ^15^. Additionally, we also used random sampling as a baseline for comparisons. Acquisition functions use either the mean (μ) and / or the 95% confidence interval width of the model output (CI_width_). The confidence interval is calculated using simulations from an ensemble of models. We distinguished three types of acquisition functions:

- Greedy: F(x) = μ(x), which considers the mean of the predictions. If a sample has a high greedy F(x) value, the corresponding perturbations are predicted to induce an upregulation of CXCL9.
- Uncertainty: F(x) = CI_width_(x), which considers the confidence interval of the model output. It represents the uncertainty of a prediction.
- GU: F(x) = μ(x) + CI_width_(x), which is a combination between the greedy and uncertainty approaches.

#### Synthetic data set

The active learning loop contains several decisions that can impact the quality of the optimization. We analyzed the effect of different configurations over multiple rounds using a synthetic data set. The synthetic data set consists of simulation of a set of conditions using the cell line-specific models. Gaussian noise was added to the values with a mean of 0 and standard deviation of 0.2 to simulate experimental noise. The synthetic data set was used to systematically assess the impact of varying the number of conditions added in each round and the selected conditions in the initial training set. When choosing the number of conditions to add, we want to create a balance between efficiency and feasibility in the wet lab. The initial training set is important as the advantage of active learning is that it is unnecessary to use all conditions from the perturbation screenings. However, it is unclear what the effect is of the initial training set on model convergence. Therefore, we investigated this effect using five different initial configurations, ranging from handpicked conditions (‘Manual’) to random sampling. For Manual, we selected conditions based on the following criteria:

1. Include condition with no perturbations (no cytokines or inhibitors)
2. Include conditions describing effects of cytokines on CXCL9 in absence of inhibitors (IFNγ, TNFα or IFNγ and TNFα only)
3. Include an inhibitor with an inhibitory effect on CXCL9 (JAKi)
4. Include one or two inhibitors with a stimulatory effect on CXCL9 (PI3Ki, MEKi)
5. Include inhibitors that had effect only in one of the two cell lines (IKKi)

#### Evaluation

Models trained with the active learning approach can be evaluated on different criteria: **(1)** the rate of detecting hits, where a hit is defined as a condition with a CXCL9 value above a specific threshold (0.2), and **(2)** the model prediction performance. For hit discovery, we valued both efficiency in detecting hits (i.e. hits are sampled in the early rounds) and correctness of predicted hits. We represented model performance for each round with the Pearson correlation coefficient (*R^2^*).

## Acknowledgements

We thank Óscar Lapuente-Santana for his guidance in setting up the prior knowledge network. This work was supported by the Netherlands Organization for Scientific Research (NWO) Gravitation program IMAGINE! (project number 24.005.009) and by NWO Aspasia grant (project number 015.021.065). ChatGPT was used to assist in language editing and improving readability. The authors reviewed and verified all generated content.

## Author contributions

B.W. carried out the wet lab experiments, implemented the computational framework and analyzed the data. V.S.M.K.Y. and L.D. supervised the project and helped design and carry out the wet lab experiments. F.E. conceived the study and was in charge of overall direction and planning. B.W. took the lead in writing the manuscript and designed the figures, with significant contribution from F.E. V.S.M.K.Y. and L.D. provided feedback on the manuscript.

## Data availability

All reagents used in this study are commercially available and suppliers are provided in the Materials and methods section. Data and additional information for perturbation experiments is provided at https://github.com/SysBioOncology/ActiveLearning_MechanisticModels_CXCL9

## Code availability

All code to ensure reproducibility of the results is available at https://github.com/SysBioOncology/ActiveLearning_MechanisticModels_CXCL9. This include preparation of the perturbation screening data, model training, active learning pipeline and parameter analyses such as *in silico* knockouts, sensitivity analysis (MPSA) and bootstrapping. Scripts to reproduce the main figures in this manuscript are also included. Installation information and package versions used for the manuscript are available in the repository.

